# Endogenous retrovirus IAP forms virus-like particles and traffics across the maternal-fetal barrier

**DOI:** 10.64898/2025.12.19.695612

**Authors:** Abby J. Bergman, Guillaume Cornelis, Julie C. Baker

## Abstract

For 50 years, virus-like particles (VLPs) have been observed in the placentas of many species, but their source and function remain unexplored. Here, we identify *intracisternal a-type particle* (IAP) elements, specifically the IAPEz-int family, as the likely source of VLPs in the mouse placenta. IAPEz-int instances are expressed throughout placentation and contain intact *gag* and *env* sequences critical for the formation of immature and mature particles, respectively. To elucidate a role for IAPEz-int derived VLPs in the mouse placenta, we generated a knock-in mouse containing a full-length element tagged with HA/FLAG. Using this line, we demonstrate that fetal IAPEz-int traffics into and across the maternal decidua. In total, we suggest that placental VLPs derive from IAPEz-int, and demonstrate their potential to traffic into maternal tissues, suggesting a role for these structures in maternal-fetal communication at the placental interface.

## Introduction

Virus-like particles (VLPs) are ubiquitous in the placentas and pregnant uterus of a variety of species, including *Homo sapiens* (human)^1–3^, *Mus musculus* (mouse)^4^, *Rattus norvegicus* (rat)^5^, *Papio anubis* (baboon)^6,7^, *Macaca mulatta* (rhesus monkey)^6^, and *Monodelphis domesticus* (opossum)^8^. Despite their pervasiveness, the genetic source of these structures has not been systematically investigated since their first observations over 50 years ago. Placental VLPs have been found across healthy samples and are morphologically consistent within, but not between, species. In *H. sapiens, P. anubis, M. mulatta, and R. norvegicus*, observed VLPs have been classified as c-type particles with a 100nm outer diameter and 50nm inner core diameter^1,2,5–7^. In *M. musculus*, the overall structure is remarkably similar, but the particles are classified as a-type and have an outer diameter of 90nm and 45nm inner core diameter^4,6^. These subtle differences in size and morphology suggest that VLPs arise from lineage specific endogenous retroviruses (ERVs), implying that convergent evolution may underlie their presence across mammals.

The presence of VLPs in the placenta raises the question of whether select ERVs escape repression in the placenta and give rise to biologically relevant functions during pregnancy. Expression of ERVs is generally detrimental as their mobilization can be mutagenic and their protein products inflammatory. Given these negative effects, ERVs are silenced by both pre- and post-transcriptional mechanisms, including DNA methylation^9–11^, histone modification^12–16^, and RNAi pathways^17^. Since the placenta is globally hypomethylated compared to other tissues^18^, ERV expression has been attributed as a non-specific result of decreased regulation. However, multiple lines of evidence, such as the presence of highly and partially methylated regions^13^ and the activity of Trim28^19^ and Piwi^20^ pathways suggest that ERVs remain faithfully silenced in the placenta. Nonetheless, the presence of VLPs alludes to a subset of the placenta ERVs that escape silencing. The select expression of ERVs, despite their devastating potential, indicates that elements that escape repression may be functionally important at the maternal-fetal interface.

VLPs arise from ERVs that have retained intact gene domains capable of producing the structural elements of a viral particle, including *gag* and *env,* whose polyproteins give rise to the capsid and envelope, respectively. In the mouse genome, the *intracisternal a-type particle* (IAP) family accounts for many of these intact domains. The ancestral IAP provirus was a stereotypical long terminal repeat (LTR) transposon, consisting of *gag, pol,* and *env* domains flanked by 5’ and 3’ LTRs^21,22^. Today, most remaining IAP instances have decayed and consist of *gag* and *pol* domains (which are variably truncated and no longer coding), flanked by LTRs^22^. *In vitro* studies of IAP have demonstrated that an intact *gag* domain is sufficient to give rise to immature a-type particles that accumulate in the endoplasmic reticulum (ER)^23^, while instances that additionally encode an *env* protein form mature particles that bud at the membrane into the extracellular space^24^. These particles have characteristics remarkably similar to placental VLPs indicating that they may derive from IAP elements with intact *gag* and/or *env* domains.

ERV-derived VLPs, such as *Arc* and *Peg10,* have been implicated in facilitating cell-cell communication by trafficking cargo between cells. In the mouse brain, Arc forms particles that traffic RNA across the synaptic junction to enable memory consolidation^25^. *In vitro*, Peg10 forms secreted VLPs which bind and stabilize a variety of mRNAs^26^. While the function of Peg10 VLPs in the placenta is unknown, deletion of *Peg10* in the mouse is embryonic lethal due to failures in proper placentation^27^. This suggests that Peg10 VLPs may have an important role in facilitating cell-cell communication via cargo delivery at the placental interface. Further, the placenta is known to shed large quantities of diverse extracellular vesicles, including VLPs, which have been implicated in a variety of functions^28^, including maternal-fetal communication^29^, immune tolerance^30,31^, and metabolism^32^. Therefore, particle-based trafficking may represent an essential facet of placental biology, enabling communication between fetal and maternal cells.

Here, we show that the ERV family IAPEz-int, is a major source of placental VLPs, is temporally and spatially restricted to the placenta, and has loci that retain intact *gag* and *env* domains. To investigate the function of IAPEz-int, we generated a transgenic mouse that contains a full-length IAPEz-int tagged with FLAG and HA. Transgenic mice display increased immature and mature VLPs in the intracellular and extracellular spaces in placental. Further, we demonstrate that fetal-derived IAPEz-int transgene is found in the maternal decidua, suggesting it traffics across the maternal-fetal interface. Finally, we report similar ERVs in other mammalian species that are selectively expressed in the placenta and retain intact *gag* domains, suggesting that VLPs may be a convergently evolved means of maternal-fetal communication. Together, these data establish IAPEz-int as a source of placental VLPs in mice and represent a novel cell-cell communication mechanism between genetically distinct tissues.

## Results

### IAP family members are expressed during mouse placentation

To determine which ERVs are expressed during placental development, we performed bulk RNAseq on mouse placentas at embryonic days (E)6.5, E9.5, and E15.5. Reads were processed and aligned using STAR^33^ with parameters that retain multimapping reads. Aligned reads were counted with TEcounts^34^ and normalized for library depth with DESeq2^35^. Critically, this approach enumerates expression levels by ERV family, which could be driven by one or many loci. To quantitatively compare the expression of each family, we assumed the null hypothesis that all elements should be equally likely to undergo spurious, unregulated transcription. Under this assumption, we expect counts to scale linearly with the total size of the family. Accordingly, we then performed linear regression between the size of the ERV family (in bp) and its normalized counts. Across gestation, we identified ten ERV families with expression levels more than three studentized residuals off the regression, representing a significant departure from the null hypothesis **(Figure 1A).** Four IAP families (IAPEz-int, IAPLTR1_Mm, IAPLTR2_Mm and IAPA_MM-int) accounted for 33% of the expressed families at E6.5 (2/6), 60% at E9.5 (3/5), and 66% at E15.5 (4/6) **(Supplemental Table 1).** IAPA_MM-int and IAPLTR1_Mm were found at all three gestational timepoints. IAPEz-int was found at E9.5 and E15.5 while IAPLTR2_Mm was only found at E15.5. The remaining six non-IAP families were restricted to single timepoints, except for MMETn-int and RLTR4_Mm which both appear at E6.5 and E9.5. This demonstrates that despite the globally hypomethylated placental environment, only select families of ERVs, predominantly IAPs, are expressed.

**Figure 1.**
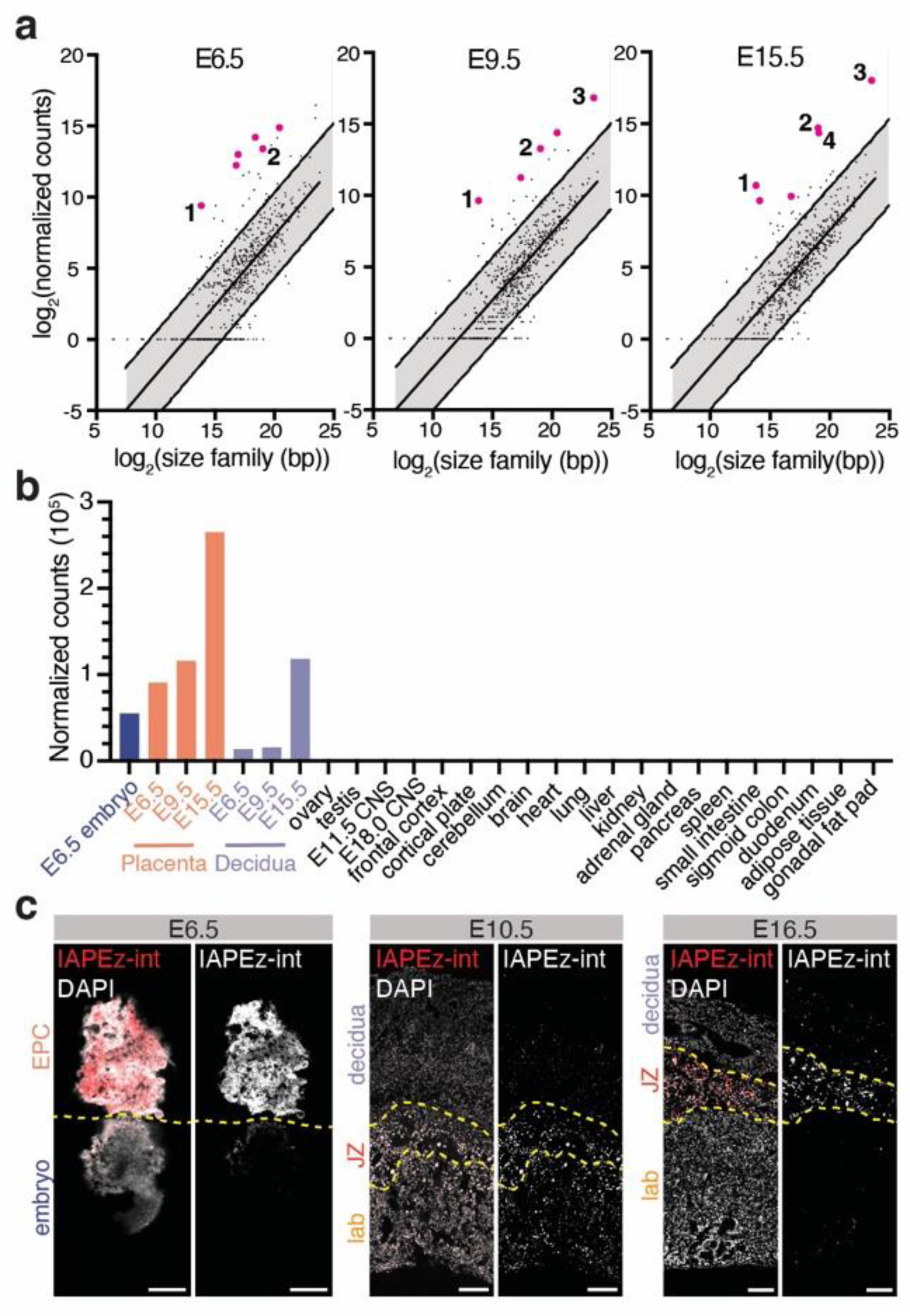
IAPEz-int is spatially and temporally expressed in the placenta. **a)** Expressed ERV families were identified at E6.5 (left), E9.5 (center), and E15.5 (right) by linear regression analysis. The line of best fit and lines at +/− 3 standard deviations are shown in solid black. The gray band encapsulates non-outliers within this error. Expressed families are plotted in magenta, with members of the IAP superfamily labeled 1-4. (1: IAPA_MM-int, 2: IAPLTR1_Mm, 3: IAPEz-int, 4: IAPLTR2_Mm). **b)** IAPEz-int expression in E6.5 embryo (blue), E6.5, E9.5, E15.5 placenta (orange) and decidua (purple), and somatic tissues of the adult and embryo (black). *CNS; central nervous system*. **c)** Maximum intensity projections of IAPEz-int *in situ* hybridization at E6.5 (left), E10.5 (center) and E16.5 (right). At each stage, the left panel shows IAPEz-int staining in red counterstained with DAPI in white and the right panel shows IAPEz-int in grayscale. *Yellow dashed lines demarcate the embryo and ectoplacental cone (EPC) at E6.5, and the labyrinth (lab), junctional zone (JZ), and decidual regions at E10.5 and E16.5. Scale bars = 200μM.*

To determine if any of the ten expressed ERV families were capable of forming VLPs, we examined these loci for *gag* domains, which are sufficient for particle formation^36^. We performed BLAST^37^ and HMMER^38^ on all ORFs belonging to an expressed ERV family. Putative *gag* domains were considered as ORFs that had sufficient homology to a retroviral *gag* domain using both approaches (*BLAST*: `e-value ≤ 1e-5, bitscore ≥ 50, pident ≥ 35, qlen ≥ 100, qcov ≥ 0.4;` *HMMER*`: score_dom ≥ 10`). In total, we identified 2511 loci containing intact *gag* domains, 2508 of which belong to the IAPEz-int family. The remaining three *gag* ORFs belonged to IAPA_MM-int (2) and MMVL30-int (1) **(Supplemental Table 2).** Additionally, we examined these loci for the presence of reverse transcriptase (*rt*) and *env* domains to assess whether they could retrotranspose and/or form mature, enveloped particles. BLAST and HMMER analysis for these domains reveal that neither the IAPA_MM-int nor MMVL30-int loci contained either *rt* or *env*. However, we identified 1279 IAPEz-int with *rt* domains and 544 with *env* domains. Further, we identified 160 IAPEz-int loci with intact ORFs for all three domains **(Supplemental Table 2)**. While we have found *gag, rt, and env* ORFs in these IAPEz-int loci, we have not investigated the individual sequences for the presence of nullifying mutations or functional regulatory elements. Regardless, we expect that some subset of these loci are capable of transposing (*rt* containing) and/or forming immature (*gag* containing) or mature (*env* containing) particles.

As IAPEz-int elements are outstanding candidates for the source of VLPs in the mouse placenta given their expression and domain homology, we next investigated their spatial and temporal expression. To determine tissue specificity of IAP families, we utilized adult and embryonic ENCODE datasets to quantify expression across tissues. We find that IAPEz-int is strongly expressed in the placenta at all stages examined, with expression increasing as gestation progresses. IAPEz-int is also expressed in the E15.5 decidua but has limited expression in other tissues **(Figure 1B).** IAPA_MM-int, IAPLTR1_Mm, and IAPLTR2_Mm display similar tissue-specific expression, while the other expressed ERV families do not **(Supplemental Figure 1).** We next performed hybridization chain reaction (HCR) *in situ* hybridization at E6.5, E10.5, and E16.5 to determine characterize IAPEz-int localization. The IAPEz-int probe set was designed to the consensus regions of published IAP sequences (92L23, 262J21^23^, MIA14^39^), tiling the *gag* and *pol* domains. At E6.5, we find IAPEz-int localized to the ectoplacental cone **(Figure 1C)**. At E10.5, we find IAPEz-int transcripts present throughout the placenta, with the highest expression in the trophoblast giant cells within the junctional zone. Interestingly, we also observe weak staining in the decidua at this time **(Figure 1C).** By E16.5, we find IAPEz-int expression to be highest in the junctional zone, with sparse, punctate signal in the decidua **(Figure 1C).** Taken together, these data demonstrate that IAP is spatially restricted expression in the developing placenta, with specific signal in the cells of the junctional zone, spreading into the maternal decidua by E16.5.

### IAP mobilization is repressed in trophoblast stem cells

Given that we identified the IAPEz-int family as being expressed in the placenta and having loci with intact *rt* domains, we examined whether these elements were capable of transposing in trophoblast stem cells. To this end, we generated a transposition reporter plasmid containing an EGFP cassette interrupted by an antisense intron^40^. This cassette was placed downstream of a transposition competent IAP element^23^, before the 3’ LTR, hereafter called (tr)IAP^EGFP^ (transposition reporter <u>IAP</u> with <u>EGFP</u>). Following transfection with (tr)IAP^EGFP^, GFP positive cells indicate the expression of a successful reverse transcription and integration event **(Figure 2A)**.

**Figure 2.**
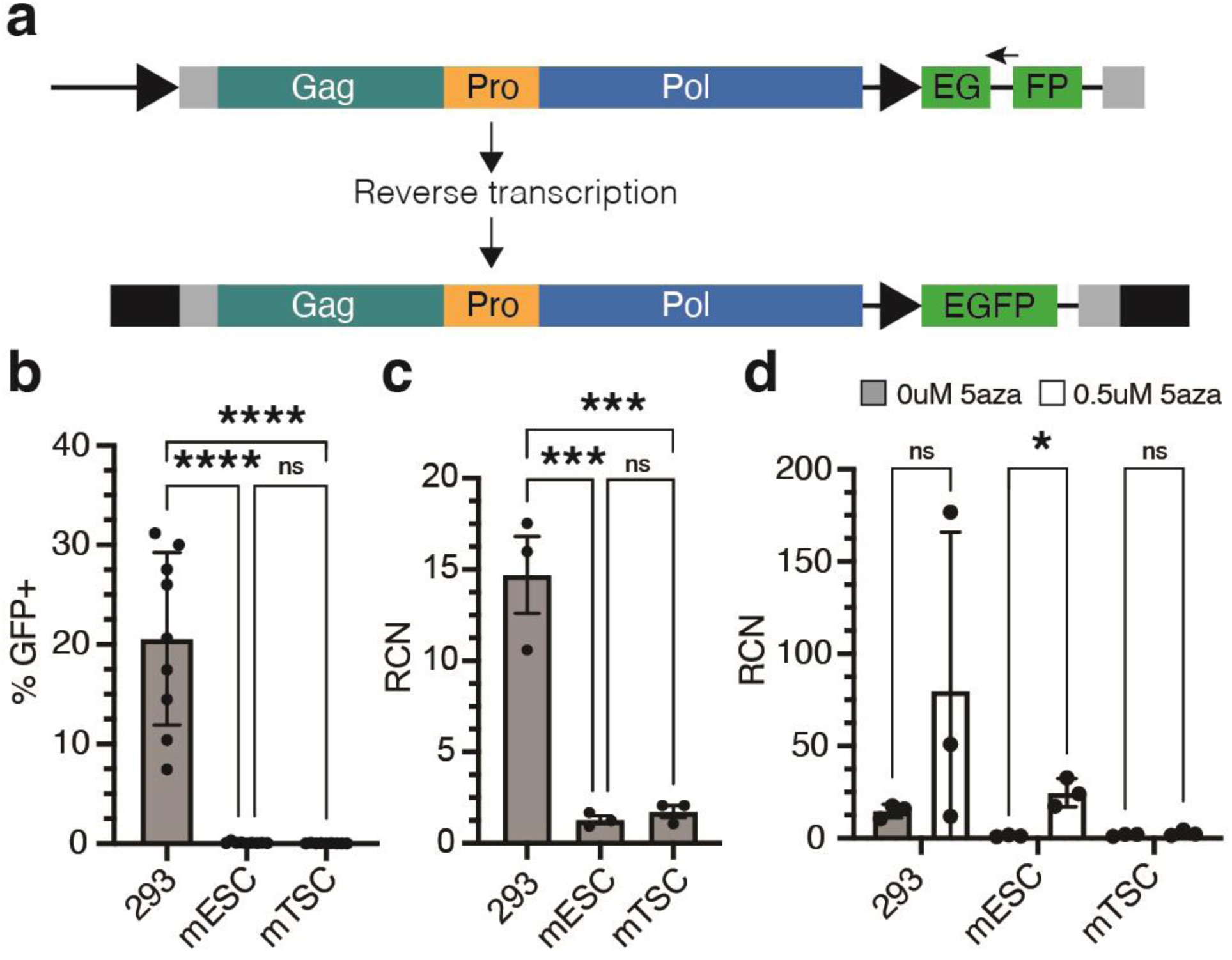
IAP transposition is repressed in embryonic lineages. **a)** Schematic of the (tr)IAP^EGFP^ transposition reporter. Teal, yellow and blue represent the *gag*, *pro*, and *pol* domains, respectively. EGFP cassette is represented in green. Right facing arrows represent CMV promoters. *Upper:* EGFP cassette with antisense β-globin intron (left facing arrow). *Lower:* Integration of the reporter following a reverse transcription event causing the splicing of the intron and reconstitution of EGFP. **b)** Quantification of the percent GFP positive cells in 293 cells, mESCs, and mTSCs after transfection with (tr)IAP^EGFP^. Statistics calculated by one way ANOVA with Tukey’s multiple comparisons correction. **c)** Relative copy number (normalized to Gapdh) of (tr)IAP^EGFP^ determined by qPCR in 293 cells, mESCs, and mTSCs. Statistics calculated by one way ANOVA with Tukey’s multiple comparisons correction. **d)** Relative copy number of (tr)IAP^EGF^ in 293 cells, mESCs, and mTSCs after 10 days with or without 5aza treatment. Gray bars represent untreated, white bars represent treated. Statistics calculated by Welch’s unpaired t-test. *RCN; relative copy number. All bars plotted represent 3-6 biological replicates. **** denotes p<0.0001, *** denotes p<0.001, * denotes p<0.05, ns: not significant.*

To determine the frequency of IAP transposition, we transfected (tr)IAP^EGFP^ into HEK293, mouse embryonic stem cells (mESCs), and mouse trophoblast stem cells (mTSCs) and calculated the transposition rate of (tr)IAP^EGFP^ normalized by the transfection rate (%GFP+ cells in (tr)IAP^EGFP^ / pEGFP). In HEK293, we determined that (tr)IAP^EGFP^ transposed in 16.50% (95% CI 9.85-23.14) of cells. In mESCs and mTSCs, (tr)IAP^EGFP^ transposed in 0.093% (95% CI 0.024-0.16) and 0.025% (95% CI 0.002-0.049) of transfected cells, respectively **(Figure 2B).** Importantly, GFP positivity in this setting indicates that both reverse transcription and integration, as well as expression of the integrated construct have occurred. Thus, the absence of GFP positivity suggests that either integration or expression of (tr)IAP^EGFP^ are inhibited in early embryonic and trophoblastic lineages.

To determine whether the integration or the expression of (tr)IAP^EGFP^ is inhibited in mESCs and mTSCs, we harvested genomic DNA from cells 10 days post transfection and quantified the relative copy number (RCN) of (tr)IAP^EGFP^ in the genome via qPCR. To discriminate between integrated copies and persistent plasmid, we amplified the EGFP cassette of (tr)IAP^EGFP^ at the splice junction, which would generate an 83bp amplicon from any integrated sequence and a 985bp amplicon from unspliced plasmid. In HEK293s, the RCN of (tr)IAP^EGFP^ was 14.7 (95% CI 5.66-23.74). In mESCs and mTSCs the RCN was 1.742 (95% CI 0.32-3.2) and 1.293 (95% CI 0.35-2.24), respectively **(Figure 2C).** This demonstrates that expression, not integration, of (tr)IAP^EGFP^ is suppressed in embryonic and trophoblast cells. Thus, we suggest that IAP is capable of reverse transcription and integration, however, its mobility is limited through silencing the expression of transposition-competent copies in these lineages.

To test whether DNA methylation silences (tr)IAP^EGFP^ expression and therefore subsequent transposition in stem cells, we treated (tr)IAP^EGFP^ transfected cells with 0.5uM 5-azacytidine (5aza) for 10 days and quantified RCN. In HEK293s, the RCN of (tr)IAP^EGFP^ was not significantly affected after 5aza treatment (p = 0.3323). In mESCs, the RCN of (tr)IAP^EGFP^ increased significantly after 5aza treatment (p = 0.0091). In contrast, in mTSCs the RCN was unaffected by treatment (p = 0.4092) **(Figure 2D).** Overall, this suggests that DNA methylation silences the expression of newly integrated IAP loci in mESCs, while an alternate mechanism is utilized in mTSCs.

### Virus-like particles with IAP characteristics are found in the mouse placenta

Given the presence of IAPEz-int loci encoding for *gag* and/or *env,* we next examined VLPs in the mouse placenta to determine if both immature and mature particles with IAP morphology could be found. To this end, we performed negative stain transmission electron microscopy (TEM) on E15.5 placentas from C3H and C57Bl/6 mice. Placentas were trimmed around the umbilical cords and fixed in 2% glutaraldehyde/4% paraformaldehyde. Samples were embedded in Epon and 75nm sections were cut *en face* to obtain grids capturing the entire umbilical-decidual face. In both strains, we identified immature VLPs displaying stereotypical, a-type morphology, with concentric, electron dense shells **(Figure 3A,B).** These immature particles were observed mainly within the ER lumen. Notably, we also observed mature particles with external envelope spikes **(Figure 3C)**. Enveloped particles were sparser but observed outside of the ER. The measured diameter of all observed particles was 66.23nm **(Figure 3D).** In total, the observed placental VLPs have features that are consistent with immature and mature IAP particles, including the size, morphology, and genetic structure. These characteristics and locations are similar to those previously published *in vitro* for IAP-derived VLPs^24^. Importantly, IAPEz-int is unique among placental expressed ERVs, having *gag* containing as well as *gag* and *env* containing loci which can give rise to both observed VLP morphologies.

**Figure 3.**
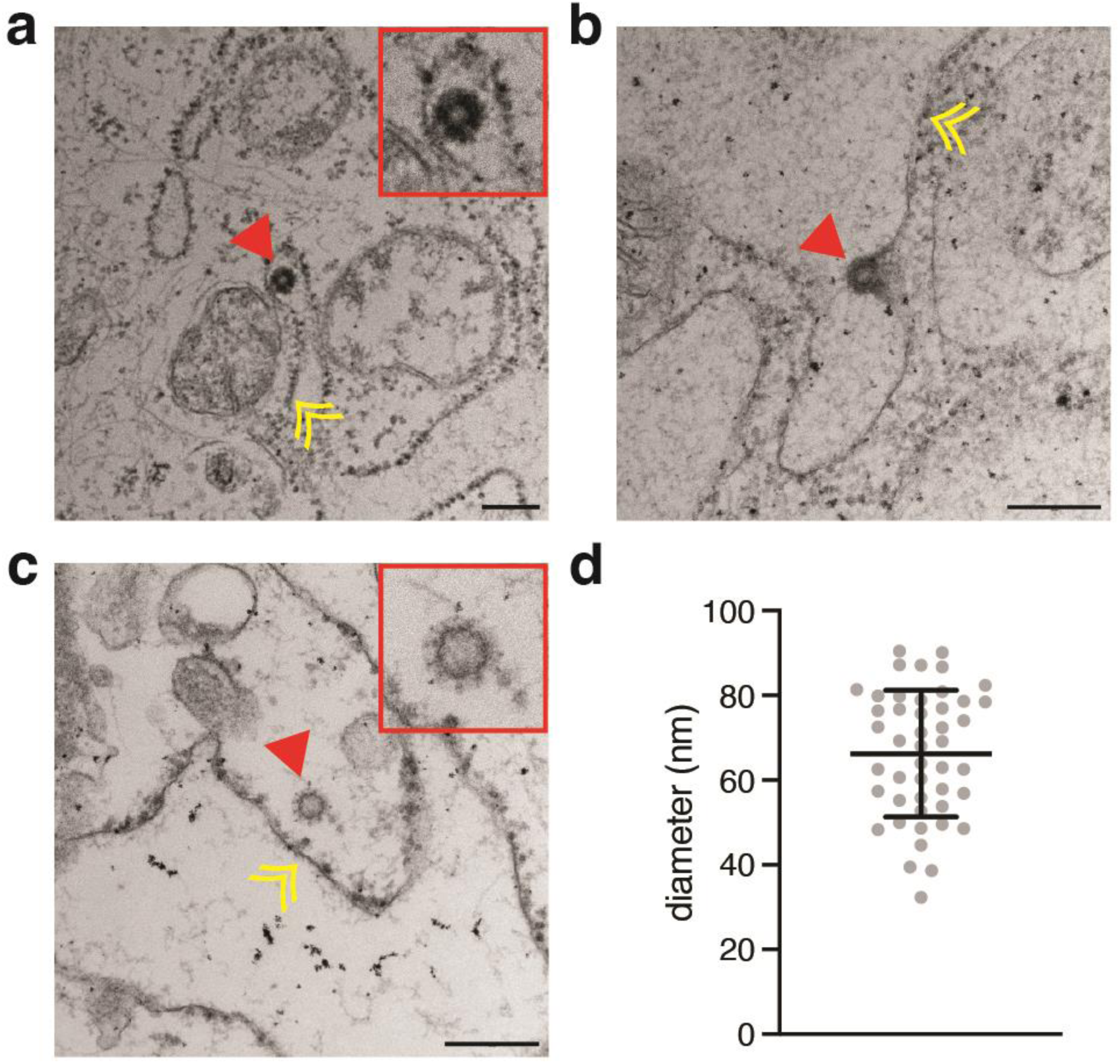
Placental VLPs have characteristics of IAP particles. **a)** Immature VLP observed in the ER. **b)** Immature VLP budding from the ER membrane. **c)** Mature VLP with spikes decorating its outer shell. **d)** Diameter of observed VLPs (n=46). Bar represents mean and standard deviation (66.23nm +/− 14.98nm). *Yellow double arrowheads denote ER membrane; red solid arrowheads denote VLPs. Red insets are high magnification of the VLPs marked by the red arrowheads. Scale bars = 100nm.*

### Enveloped IAP is secreted extracellularly

We next investigated the subcellular localization of putative mature (*gag* and *env* containing) and immature IAPEz-int (*gag* only) VLPs *in vitro*. To this end, we generated two expression plasmids, one consisting of only the IAPEz-int *gag* domain (pIAP*gag*^HA^), and the other consisting of IAPEz-int *gag, pol,* and *env* (pIAP*env*^FLAG/HA^). Both constructs were tagged with HA at the C terminus **(Figure 4A)**. HEK293 cells were transfected with either expression plasmid and the supernatant and lysate were harvested two days post-transfection. We then performed western blotting with an anti-HA antibody to detect the presence of IAP in the cellular or extracellular fractions. In the cell lysate, we detect both IAP*gag* and IAP*env* **(Figure 4B),** while only IAP*env* is present in the supernatant **(Figure 4C).** This is consistent with our TEM observations and others’ ^24^, and demonstrates that the *gag* domain of IAP alone is not sufficient for extracellular secretion.

**Figure 4.**
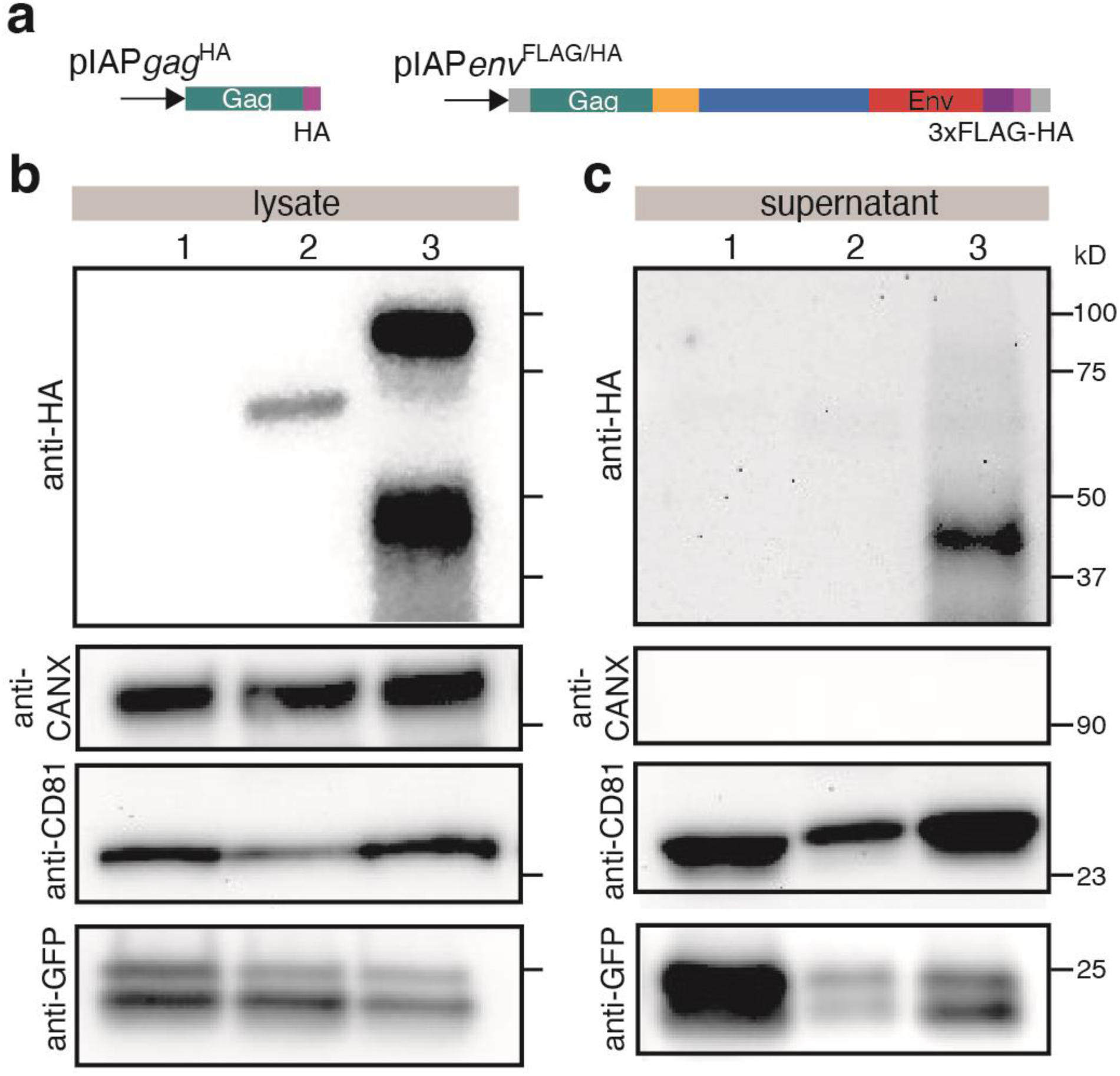
Cellular behavior of IAP is dependent on intact gene domains. **a)** Schematics of pIAP*gag*^HA^ (left) and pIAP*env*^FLAG/HA^ (right) plasmids. Teal – *gag;* yellow – *pro;* blue – *pol*; red – *env;* purple – HA, 3xFLAG/HA. **b)** Western blots of HEK293 cellular lysates probed with anti-HA, anti-CANX, anti-CD81 or anti-GFP antibodies. **c)** Western blots of supernatants probed with anti-HA, anti-CANX, anti-CD81 or anti-GFP antibodies. *Lane 1:* pEGFP *transfected. Lane 2:* pIAP*gag*^HA^ *&* pEGFP *transfected. Lane 3:* pIAP*env*^FLAG/HA^ *&* pEGFP *transfected.*

### IAP overexpression leads to an increase in placental VLPs

To test whether IAP is sufficient to produce VLPs in the placenta, we generated a targeted knock-in mouse with an HA/FLAG tagged, full length IAP element at the Rosa26 locus (B6-Gt(ROSA)26Sor^tm1(CAG-IAPED1-3xFLAG-HA)^, hereafter R26-IAP). The construct consisted of the IAPE-D1 proviral sequence^24^, including the flanking LTRs, tagged C-terminally with 3xFLAG and HA. We used integration mediated transgenesis^41^ to insert this sequence (*CAG-IAPED1-3xFLAG-HA*, hereafter *Iap(fh)*) at the *Gt(ROSA)26* locus under a ubiquitous CAG promoter. R26-IAP animals display no obvious phenotypic abnormalities, and the locus is inherited in a Mendelian fashion **(Supplemental Figure 2)**. To characterize the expression of *Iap(fh)*, we performed qPCR, western blotting, and *in situ* hybridization. We first examined the transcription of *Iap(fh)*. To this end, we amplified *Iap(fh)* transcripts specifically with primers binding in the HA tag and internal *env* sequence. We observed that *Iap(fh)* is highly expressed in the placenta, and that its presence only moderately increases endogenous IAPEz-int expression **(Figure 5A).** To determine whether *Iap(fh)* is translated into protein, we harvested total protein from E9.5 and E15.5 extraembryonic and embryonic tissues (whole embryo (E9.5) or brain, lung, and kidney (E15.5)) and performed western blotting with an anti-FLAG antibody. We find that FLAG is present in all tissues of transgenic animals, indicating the presence of Iap(fh) protein **(Figure 5B)**. Next, we performed *in situ* hybridization with endogenous IAPEz-int and *Iap(fh)* probe sets in the E15.5 placenta. We show that *Iap(fh)* transcripts are present throughout the entire placenta, while endogenous IAPEz-int transcripts remain largely restricted to the junctional zone, as in wildtype **(Figure 5C).** Together, these data demonstrate that *Iap(fh)* is ubiquitously expressed and has no effect on maternal or fetal viability.

**Figure 5.**
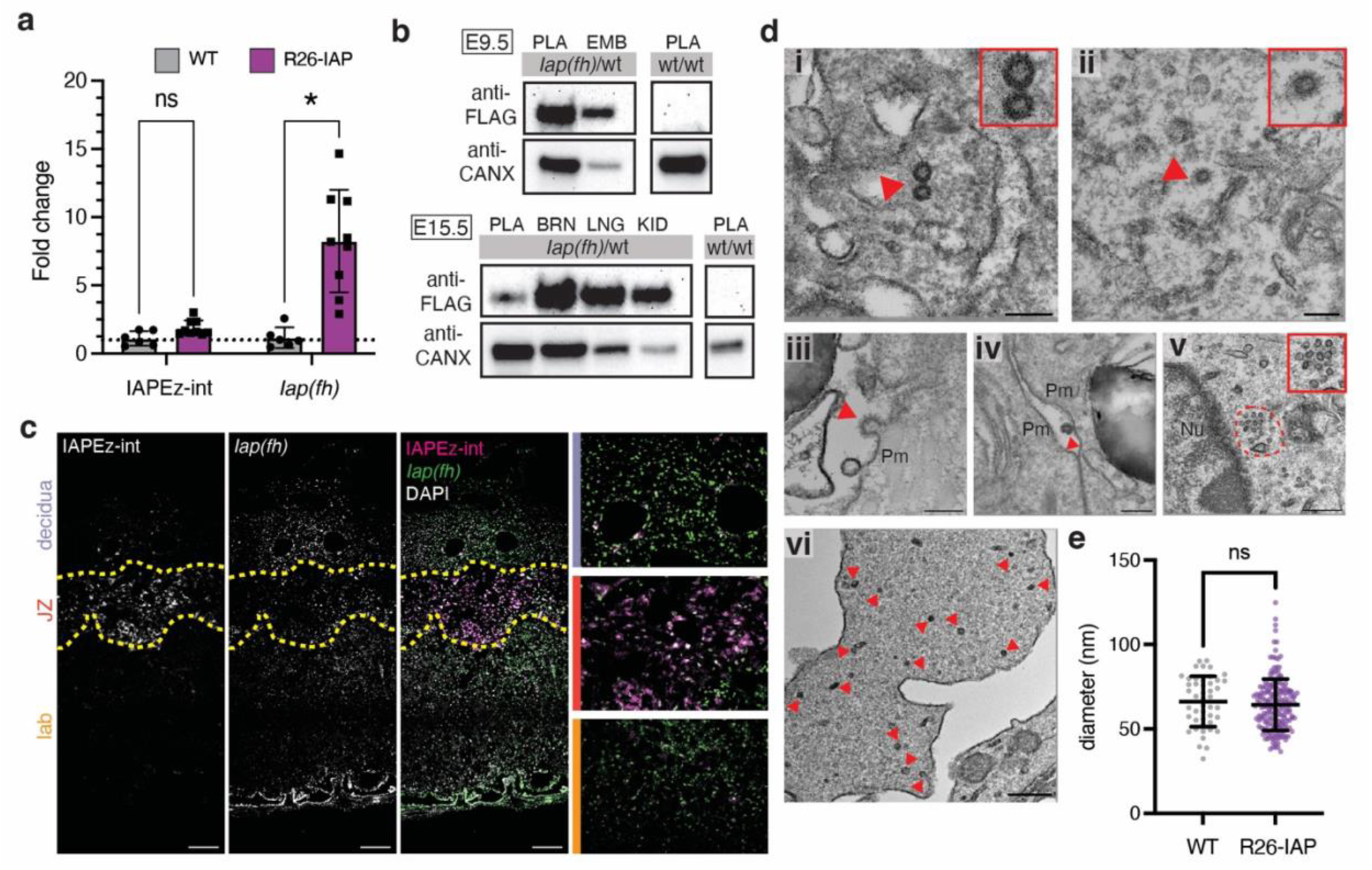
IAP knock-in mice express Iap(fh) and form copious placental VLPs. **a)** qPCR of endogenous IAPEz-int and *Iap(fh)* in wildtype and R26-IAP mouse placentas normalized to wildtype expression. Gray bars represent wildtype samples (n=2 animals from 2 litters), purple bars represent R26-IAP samples (n=3 animals from 1 litter). All measurements are averages of technical triplicates. Dashed line denotes fold change = 1. Statistics calculated by Welch’s unpaired t-test. **b)** Western blot using anti-FLAG and anti-CANX antibodies. *Upper:* E9.5 placenta (PLA) and whole embryo (EMB). *Lower:* E15.5 placenta (PLA), brain (BRN), lung (LNG), kidney (KID). Gray boxes state *Iap(fh)* genotype. **c)** Maximum intensity projection of multiplexed *in situ* hybridization of E15.5 R26-IAP placenta. The left two panels show IAPEz-int and *Iap(fh)* staining in grayscale. The right two panels show composite staining with IAPEz-int in magenta, *Iap(fh)* in green, and DAPI in white. *Yellow dashed lines demarcate the decidua, junctional zone (JZ), and labyrinth (lab).* Rightmost images are insets of (top to bottom) the decidua, junctional zone, and labyrinth. *Scale bars = 200μM.* **d)** VLPs observed in TEM of E15.5 R26-IAP placentas. **i)** Immature and **ii)** mature particles are observed in the cytoplasm of cells. **iii)** Mature particle budding from the plasma membrane. **iv)** Particle in the extracellular space. **v)** Particles are observed in high density patches. **vi)** Numerous particles observed in an extracellular vesicle. *Pm: plasma membrane, Nu: nucleus. Red arrowheads and dashed lines denote VLPs. Red boxes are insets of VLPs at red annotations. Scale bars = 100nm (i-iv), 200nm (v-vi).* **e)** Quantification of observed VLP diameter in wildtype (gray, n=46) and R26-IAP (purple, n=191) placentas. Bars represent mean and standard deviation (wildtype: 66.23nm +/-14.98nm, R26-IAP: 64.36nm +/− 15.26nm). Statistics calculated by two-tailed Mann-Whitney test. *\* denotes p<0.05, ns = not significant.*

To investigate the frequency and morphology of VLPs in R26-IAP placentas, we performed TEM on E15.5 placentas as before. Throughout these sections, we observed immature **(Figure 5D,i)** and mature **(Figure 5D, ii)** VLPs with concentric, electron dense shells and an average diameter of 64.36nm **(Figure 5E)**. VLPs observed in R26-IAP placentas were found ubiquitously throughout the cell and in extracellular features **(Figure 5D, iii-iv)**. Importantly, VLPs in these placentas are highly abundant and often found in patches of 5 or more **(Figure 5D, v-vi)**. This demonstrates that Iap(fh) efficiently forms VLPs in the placenta, and that these particles retain wildtype IAP VLP morphology. The increase in number and variety of particles in R26-IAP placentas establishes this mouse as a model to investigate the function of placental VLPs.

### *Iap(fh)* traffics across the placental barrier into maternal uterine cells

We next examined whether *Iap(fh)* could traffic across the maternal-fetal barrier. To this end, wildtype females were crossed with R26-IAP males. In resulting pregnancies, *Iap(fh)* is genetically restricted and expressed only in fetal tissues (including the fetal placenta) as the mother does not carry the transgene. We collected placentas at E9.5 and E15.5 from these crosses and performed *in situ* hybridization for *Iap(fh)*. At E9.5, we observed punctate *Iap(fh)* signal in the decidua adjacent to the placental trophoblast giant cells **(Figure 6A**, arrowheads). At E15.5, we find broad *Iap(fh)* signal throughout the maternal decidua **(Figure 6B).** This suggests that *Iap(fh)* is transported from the placenta into the decidua, even as far as the myometrial boundary.

**Figure 6.**
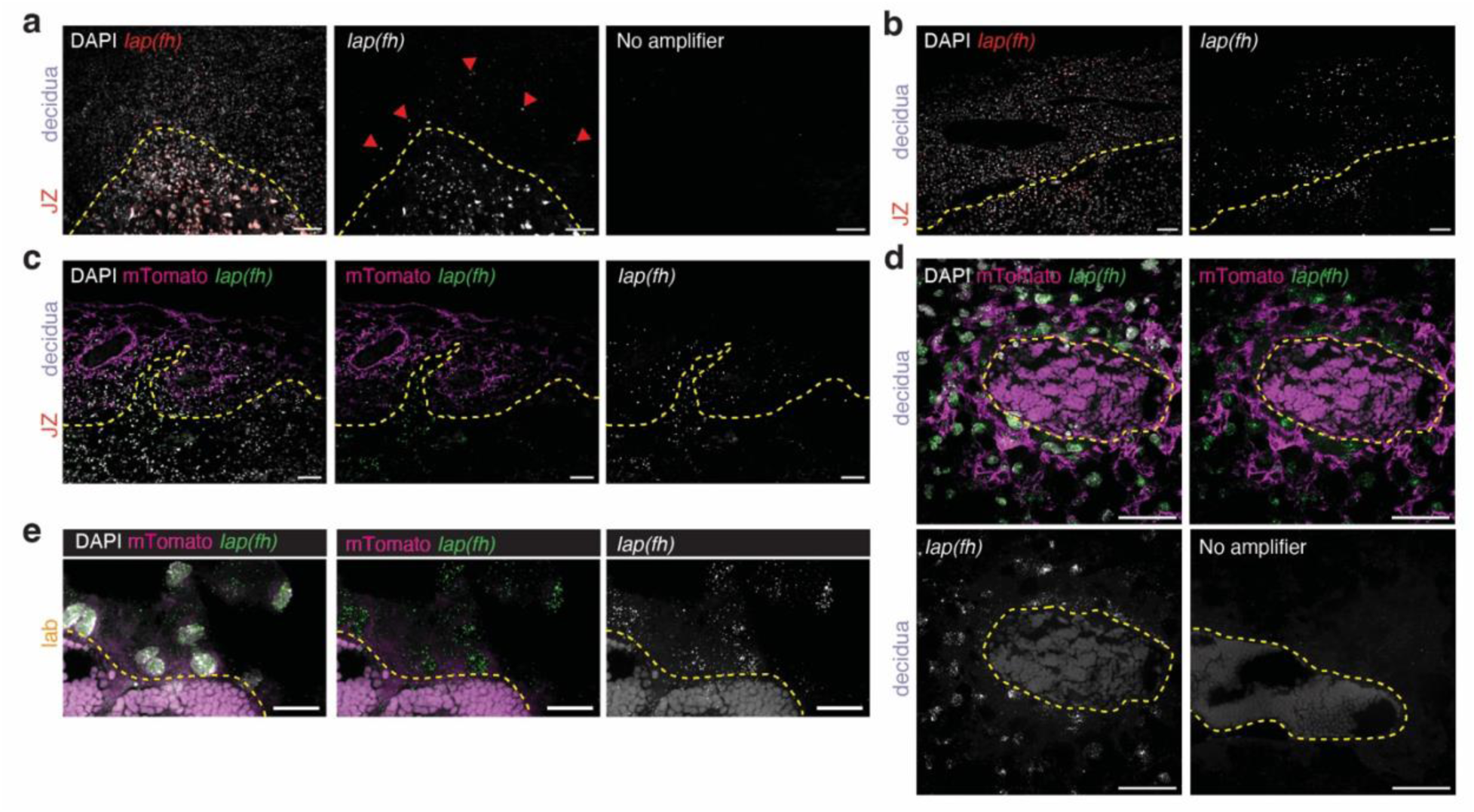
. **Iap(fh) traffics across the placental barrier into maternal tissue a)** *In situ* hybridization of *Iap(fh)* in the placenta at E9.5. Panels (left to right) show *Iap(fh)* signal (red) counterstained with DAPI (white); *Iap(fh)* signal in grayscale; and no amplifier control*. Red arrowheads denote Iap(fh) puncta in the maternal decidua.* **b)** *In situ* hybridization of *Iap(fh)* in the placenta at E15.5. Left panel shows *Iap(fh)* signal (red) counterstained with DAPI (white). Right panel shows *Iap(fh)* signal in grayscale. **c)** *In situ* hybridization of *Iap(fh)* in the placenta at E15.5 in an *mTmG*-positive; *Iap(fh)-*negative maternal background. The left panel shows the composite of *Iap(fh)* staining (green), mTomato (magenta), and DAPI (white). Center panel shows the composite of *Iap(fh)* (green) and mTomato (magenta). The right panel shows *Iap(fh)* in grayscale. **d)** High magnification of a maternal blood space from **(c).** The top left panel shows the composite of *Iap(fh)* (green), mTomato (magenta), and DAPI (white). The top right panel shows the composite of *Iap(fh)* (green) and mTomato (magenta). The bottom left panel shows *Iap(fh)* in grayscale. The bottom right panel shows the no amplifier control. **e)** *In situ* hybridization of *Iap(fh)* in the placental labyrinth at E15.5 in an *mTmG*-positive; *Iap(fh)-*negative maternal background. The left panel shows the composite of *Iap(fh)* (green), mTomato (magenta), and DAPI (white). The center panel shows the composite of *Iap(fh)* (green) and mTomato (magenta). The right panel shows *Iap(fh)* signal in grayscale. *Yellow dashed lines demarcate the decidua, junctional zone (JZ), and labyrinth (lab) **(a-c)** or the blood lumen **(d-e)**. Scale bar = 100μM **(a-c)**, 50μM **(d)**, 20μM **(e)**.*

We then asked if we could identify maternal cells encapsulating *Iap(fh)* from the fetus. To this end, we utilized R26-mTmG females which constitutively express membrane Tomato (mTomato) in all cells. These animals were bred to R26-IAP males, giving rise to pregnancies where *Iap(fh)* is only transcribed in placental cells and mTomato marks only maternal cells. We harvested E15.5 placentas from these crosses and performed *in situ* hybridization for *Iap(fh)* **(Figure 6C).** Remarkably, we found that *Iap(fh)* was contained within cells marked by mTomato throughout the decidua **(Figure 6C)**. *Iap(fh)* signal was concentrated in cells surrounding maternal blood sinuses **(Figure 6D)**, which are the passageway between the mother and fetus. To understand if *Iap(fh)* is being shunted into the bloodstream to be trafficked into the decidua, we examined *Iap(fh)* in the placental labyrinth. Indeed, we observed *Iap(fh)* signal in the fetal cells surrounding blood spaces **(Figure 6E).** Taken together, these data suggest that *Iap(fh)* may be shed by the fetal cells into the blood stream where it then travels to maternal tissue.

### VLP-competent ERV expression is present in placentas of many species

We next asked if ERVs with VLP-competent *gag* sequences are expressed in species spanning variable placental morphologies and classifications from across *Eutheria* and *Metatheria*. To identify expressed ERVs, we performed regression analysis on RNAseq data from the placentas of 12 species (see **Supplemental File 1** for source publications and accessions). BLAST and HMMER were then used to identify ORFs encoding *gag* within these expressed families. With the exception of *Loxodonta africana* (African elephant), we identified at least one family that is expressed in the placenta in all examined species **(Table 1, Supplemental Table 3)**. Homology searches reveal intact *gag* ORFs among the expressed ERVs in 6 of the 12 species. These include species where VLPs have previously been observed (*H. sapiens*^1–3^, *M. mulatta*^6^*, R. norvegicus*^5^*, M. domesticus*^8^), and 2 additional species (*Equus caballus, Sus scrofa)* **(Table 1, Supplemental Table 3).** Despite the presence of expressed ERVs, we did not identify intact *gag* domains in the 5 remaining species (*Bos taurus, Ovis aries, Canis familiaris, Dasypus novemcinctus, Pan paniscus*). This is likely due to the lack of numerous quality datasets that span total gestation time in these species. Regardless, these data suggest that the presence of placental VLPs is widespread across Mammalia, and that these VLPs may facilitate fundamental placental processes.

**Table 1.** ERV families with gag ORFs are expressed in numerous placental species.

| **Species**<br>**(common**<br>**name)** | **Infraclass** | **Placental**<br>**morphology** | **Trophoblast**<br>**invasiveness** | **Expressed**<br>**families** | **Candidate**<br>**VLP loci** |
| --- | --- | --- | --- | --- | --- |
| *Bos taurus*<br>(cow) | Eutheria | Cotyledonary | Synepitheliochorial | 1 | 0 |
| *Ovis aries*<br>(sheep) | Eutheria | Cotyledonary | Synepitheliochorial | 9 | 0 |
| *Canis familiaris*<br>(dog) | Eutheria | Zonary | Endotheliochorial | 3 | 0 |
| *Loxodonta*<br>*africana*<br>(African<br>elephant) | Eutheria | Zonary | Endotheliochorial | 0 | 0 |
| *Dasypus*<br>*novemcinctus*<br>(9-banded<br>armadillo) | Eutheria | Zonary /<br>discoid | Hemochorial | 1 | 0 |
| *Pan paniscus*<br>(bonobo) | Eutheria | Discoid | Hemochorial | 1 | 0 |
| *Macaca mulatta*<br>(rhesus<br>macaque) | Eutheria | Discoid | Hemochorial | 29 | 326 |
| *Rattus*<br>*norvegicus* (rat) | Eutheria | Discoid | Hemochorial | 21 | 718 |
| *Homo sapiens*<br>(human) | Eutheria | Discoid | Hemochorial | 51 | 44 |
| *Mus musculus*<br>(mouse) | Eutheria | Discoid | Hemochorial | 6 | 2509 |
| *Sus scrofa* (pig) | Eutheria | Diffuse | Epitheliochorial | 2 | 64 |
| *Equus caballus*<br>(horse) | Eutheria | Diffuse | Epitheliochorial | 7 | 62 |
| *Monodelphis*<br>*domesticus*<br>(opossum) | Metatheria | Yolk sac | Epitheliochorial | 1 | 77 |

## Discussion

The presence of VLPs in the placenta has been reported for many decades, yet their source and function has remained a mystery. Here, we suggest that the source of these structures in mice are IAPEz-int loci which specifically escape repressive mechanisms in the placenta. IAPEz-int is the only ERV whose expression is restricted to the placenta throughout gestation, and which contains the intact gene domains to account for the observed VLPs. Further, in the placentas of transgenic mice constitutively expressing an IAP element, we observe clustered, abundant immature and mature VLPs. When we selectively restrict the expression of this transgene to the fetus, we observe its signal in the maternal decidua, demonstrating that IAPEz-int can traffic directionally across the fetal-maternal boundary. Further, we demonstrate that similar ERVs can be identified in many mammalian species, suggesting that the function of placental VLPs is not mouse specific, but another convergently evolved trait of placentation.

### The placenta retains mechanisms to silence ERV transcription

In this study, we reveal that most ERVs are silenced in the placenta with only a handful of families, chiefly IAPs, evading repression. Given that the placenta is globally hypomethylated, this suggests that most ERVs are silenced by other mechanisms. This is consistent with what is known in mESCs and human neural progenitor cells (NPCs), which employ histone modification and post-transcriptional pathways to ensure robust silencing of ERV^10,14–16,42–44^. Thus, the question arises of how particular IAP loci escape repression in the placenta. Unfortunately, there is a dearth of data from the placenta to draw on to answer this question, especially across gestation. However, the silencing mechanisms of the IAP family have been studied in embryonic and adult cells, revealing substantial variety in the regulatory mechanism employed depending upon the developmental and cellular context. IAP loci are silenced via DNA methylation in male germ cells^45^ and in the E8.5 embryo^11^; by H3K9me3 in primordial germ cells^46^, visceral endoderm^47^, and NPCs^42^; and by DICER in the pre-implantation embryo^17^ and mESCs^44^, demonstrating that DNA methylation is not the sole mechanism by which IAPs can be silenced. In total, our results challenge the idea that genome hypomethylation drives a broadly permissive environment for ERV expression in the placenta. Rather, alternate mechanisms, including histone modification, remain active and robust in silencing TEs.

While the repressive mechanism of an ERV can vary by cellular context, the age of the element also appears to correlate with silencing programs. Jönsson and colleagues demonstrated that the age of an element dictates the dominant repressive mechanism, with the younger elements specifically subject to TRIM28-mediated H3K9me3 silencing^10^. IAP is a prolific retrotransposon, having numerous copies that have retained transposition-competency to generate new integrations in the mouse genome^21,23^. These new integrations are therefore likely to be predominantly repressed by Trim28. Interestingly, Enriquez-Gasca, *et al* have demonstrated that evasion of H3K9me3 silencing occurs through mutation of the TRIM28 binding motif in the IAP LTR^48^, suggesting that the expressed IAPEz-int instances in the placenta contain this mutation. As younger elements are more likely to be transposition competent, this mutation would be costly to their mobility. Taken together, this suggests that IAPEz-int loci that are expressed in the placenta cannot transpose, allowing the tissue to safely leverage their VLP-forming ability without the threat of their mutagenic potential.

### IAP-derived VLPs may modulate maternal-fetal immune tolerance

We propose that the function of IAP-derived VLPs is to mediate immune tolerance of the fetus by trafficking into the maternal decidua. This could be carried out by one or both of two distinct mechanisms – first, by trafficking RNA cargo, and second, by acting by itself as an antigen to uterine mast cells. VLP-based intercellular communication has strong precedence, as illustrated by Arc, which binds RNAs and traffics them between neurons^25^. While VLPs can package a variety of cargos, binding of single stranded RNA is a native function of these proteins^49^. Delivery of ssRNA into the maternal decidua could have many roles, including modulating a pro-inflammatory response to mediate immune tolerance^50,51^. Furthermore, the IAP protein itself, independently of its conformation as a VLP, is predicted to be an IgE-binding like protein, and in this capacity, may act as an antigen to modulate the decidual environment. IAP binding of IgE on uterine mast cells could lead to the degranulation of mast cells and/or activation of the COX-2 pathway. In turn, this would modulate vascular permeability, or increase prostaglandin synthesis, respectively^52^. In total, we propose a putative function for VLPs in the placenta as a mechanism to mediate and modulate the maternal-fetal relationship. This data reframes the presence of placental VLPs as merely an unusual, sporadic observation to a bona fide, functional aspect of placental biology.

### Convergent evolution of VLP mediated cell-cell communication

Convergent evolution is a strategy adopted to facilitate placental function across species. This is spectacularly illustrated by the repeated emergence of the placenta in species as diverse as fish, lizards, mammals, and arthropods. In addition, the genetic and molecular mechanisms that facilitate the placenta’s core functions also appear to have been independently assembled using similar but species-specific tools. This suggests that the maternal-fetal interface applies a strong selective pressure for the cellular and molecular strategies at play to enable successful viviparity. One of the most fantastic examples of this convergent evolution is the placental *syncytins.* Syncytins arise from the *env* sequence of ERVs, and when expressed in the placenta, facilitate the fusion of trophoblasts to generate the multinucleate syncytiotrophoblast^53,54^. These syncytia are critical for placental function and fetal survival^55,56^. Interestingly, the *env* genes that gave rise to *syncytins* are distinct in each lineage, having originated from species-specific ERVs^53,54,57^. Yet, this evolution has been observed across Therians, and even in the Mayuba lizard^57^. There are many further examples of convergent evolution in the placenta, including species-specific suites of pregnancy hormones^58^, the recruitment of amino acid transporters for nutrient transfer^59^, and prolactin-driven nutrient provisioning in mammary glands^60^, placentas^61^, and maternal follicles^62^. The pressure of the Red Queen race^63^ at the maternal-fetal interface has been proposed to be so great that evolutionary change encompasses many new strategies, including convergent evolution. In essence, there seem to be limited tools with which this evolutionary pressure can be addressed, leading to independent and repeated solutions, even among diverse species.

We would like to consider the possibility that placental VLPs are not an artifact of stray ERV expression, and are rather one of these repeatedly evolved solutions, used to mediate cellular communication across the maternal-fetal interface. VLPs were first reported at the placental interface of numerous mammals over 50 years ago, yet their source and function remain elusive. We show that ERVs encoding intact *gag*s are expressed in the placentas of many species, spanning diverse morphologies, suggesting that VLPs may be a converged feature of placental biology. In the mouse, we identify IAPEz-int as the source of placental VLPs, as it (1) is highly and specifically expressed in across gestation (2) retains intact *gag* and *env* domains, and (3) travels across the placental interface into the decidua. IAP elements are specific to the rodent lineage and therefore cannot account for the presence of VLPs in other species. Thus, VLPs in each species are likely to be derived from lineage specific ERVs, suggesting a mechanism of convergent evolution. Further characterization of the source and function of VLPs across placental morphologies has the potential to reveal physiological aspects of the maternal/fetal interface that drive a new and innovative form of cellular communication.

## Methods

### Mice

All animal husbandry and experiments were carried out in accordance with APLAC and institutional guidelines set forth by the VSC at Stanford University. C57Bl6/J, C3H, and mTmG (B6.129(Cg)-Gt(ROSA)26Sor^tm4(ACTB-tdTomato,-EGFP)Luo^/J) mice were obtained from Jackson Laboratories. For timed embryo dissections, copulation was determined by the presence of a vaginal plug the morning after mating and timings were established with embryonic day E0.5 being noon of that day. Tissues were dissected following sacrifice of the dame on the designated embryonic day. For embryos and placentas, conceptuses were removed from the uterus and separated from the decidua before separating by severing the umbilical cord. For decidual samples, tissues were cut at the antimesometrial side to limit placental contamination (E6.5, E9.5) or peeled off the placental disc (E15.5). Tissues were processed as follows:

#### OCT

Tissues were fixed in 4% PFA (VWR 100503-917) overnight at 4C. The following morning, tissues were washed 3X in PBS, then equilibrated in 30% sucrose overnight at 4C. After tissues were equilibrated (sank), they were mounted in disposable molds and frozen at −20C in OCT (Fisher Scientific 23-730-571). 8uM sections were cut on a Leica cryostat, mounted on SuperFrost Plus slides (Fisher Scientific 12-550-15), and stored at −80C until use.

#### Wholemounts for in situ hybridization

E6.5 tissues were fixed in 4% PFA for 10 minutes at room temperature and washed 3X in PBS on ice. Tissues were stored up to 1 week in PBS before use. E9.5 and later tissues were fixed in 4% PFA overnight at 4C. The following morning, tissues were washed 3X in PBS, then 2X in PBST (0.1% Tween) on ice. Tissues were then dehydrated through a graded methanol series (25% MeOH in PBST, 50%, 75%, 100%) and stored in 100% methanol at −20C for up to 6 months.

#### Total RNA extraction

Tissues were homogenized with a rotor-stater in Trizol reagent (ThermoFisher Scientific 15596026) and stored at −20C until use.

#### Protein extraction

Tissues were minced with a razor blade, snap frozen in LN_2_, and stored at −80C until use.

### Plasmids and cloning

pEGFP (Addgene Plasmid #36412) was a generous gift from Chris Kaelin.

#### pIAPgag^HA^

HA tag was added C-terminally to the IAPEz-int *gag* ORF consensus sequence. The resulting sequence was inserted downstream of the EF1a promoter into the pTwist-EF1a backbone. The resulting plasmid was synthesized by Twist Bioscience (South San Francisco, California, USA).

#### (tr)IAP^EGFP^

Transposition competent IAP92L23 sequence was retrieved from GenBank (AC012381 pos 161601-168684, + strand). The split EGFP transposition cassette was identified from the pWA125 vector and consists of an independent promoter, Kozak sequence, EGFP coding sequence with intervening antisense b-globin intron, and independent polyA signal. To generate the (tr)IAP^EGFP^ construct, the EGFP cassette was inserted antisense to IAP ORFs downstream of the Pol domain (pos 5923 in IAP92L23). The resulting sequence was synthesized in 3 fragments (fragment 1 pos 1-4762, fragment 2 pos 4763-7038, fragment 3 pos 7038-9583) by Invitrogen (Waltham, Massachusetts, USA). pcDNA3.4 mammalian expression vector was linearized with KpnI and NotI and fragments 1-3 were PCR amplified using NEB Q5 HiFi polymerase (New England Biolabs M0491) and assembled with NEB HiFi mastermix (New England Biolabs E2621). The final plasmid then contains the transposition competent IAP92L23 with split EGFP reporter under a CMV promoter and a WPRE and TK polyA signal.

#### pIAPenv^FLAG/HA^

The full length IAPE-D1 proviral sequence was retrieved from GenBank (AC123738, pos 116181-152862, - strand). The sequence was then modified to include a 3xFLAG + 5xGly spacer + HA tag immediately preceding the env stop codon (pos 7578 in IAPE-D1 sequence). The 5’ sequence was truncated to begin at the R-U5 of the 5’LTR (pos 226 of the IAPE-D1 sequence). The resulting sequence was synthesized in 2 fragments (fragment 1 pos 1-3478, fragment 2 pos 3479-8225) by Invitrogen (Waltham, Massachusetts, USA). Fragment 1 was synthesized in the mammalian expression vector pcDNA3.4. Fragment 1 was linearized with KpnI and NotI and fragment 2 was PCR amplified using NEB Q5 HiFi polymerase and then assembled with NEB HiFi mastermix. The final plasmid then contains the tagged, IAPE-D1 sequence under a CMV promoter and a WPRE and TK polyA signal following the IAPE-D1 3’LTR.

#### pBT378_IAPenv^FLAG/HA^

To generate the plasmid used to inject into landing pad mice, the IAPE-D1(FLAG/HA) sequence (as described above) was cloned into the pBT378Linker plasmid (gift from Stanford Transgenic, Knockout, and Tumor Model Center). The pBT378Linker backbone was linearized with SbfI and NruI. IAPE-D1(FLAG/HA) was PCR amplified in two fragments (fragment 1 pos 1-3478, fragment 2 pos3479-8225) using NEB Q5 HiFi polymerase and assembled with NEB HiFi mastermix. The final plasmid then contains the tagged, IAPE-D1 sequence under a CAG promoter and a 3’ SV40 polyA element following the 3’LTR. This entire construct is contained between attB sites in the backbone which are utilized to facilitate the integration into the landing pad mouse.

### Tissue Culture

HEK293 cells (ATCC CRL-1573) were maintained in DMEM (Thermo Scientific 11995065) supplemented with 10% v/v FBS (Cytiva SH30071.03), 2mM glutaMAX (ThermoFisher Scientific 35050061), 100U/mL penicillin, 100ug/mL streptomycin (ThermoFisher Scientific 15-140-122), and 1mM sodium pyruvate (Sigma-Aldrich S8636). To introduce plasmid DNA, 293 cells were transfected using Lipofectamine 2000 reagent (ThermoFisher Scientific 11668027). All transfections were conducted according to the manufacturers protocol with the following parameters per well for a 12-well format: seeding density: 100,000 cells, 2ug plasmid DNA, 2.5uL Lipofectamine reagent. For other plate formats, parameters were scaled according to surface area.

mTSCs were maintained in DMEM/F-12 (ThermoFisher Scientific 11330032) supplemented with 20% v/v FBS, 2mM glutaMAX (ThermoFisher Scientific 35050061), 100U/mL penicillin, 100ug/mL streptomycin, 1mM sodium pyruvate, and 50uM 2-mercaptoethanol (Fisher Scientific 21-985-023) supplemented with 25ng/mL Fgf4 (R&D Systems 235-F4-025), 1ug/mL ActivinA (R&D 338-AC-050), and 10ng/mL Heparin (Sigma-Aldrich H3149-10KU). To introduce plasmid DNA, mTSCs were transfected using the NEON electroporation system (Thermo Scientific MPK10025). All transfections were conducted according to the manufacturers protocol with the following parameters for a 100uL tip: 1 x 10^6^ cells / 100uL, 10ug plasmid DNA. Electroporation parameters were as follows: 1125V, 20ms, 2 pulses.

mESCs were maintained in the absence of feeder cells on plates coated with laminin (Thermo Scientific 23017015) and poly-L-ornithine (Sigma-Aldrich P4638-100mg). mESCs were maintained in 2i medium composed of 50% DMEM, 50% Neurobasal (Thermo Scientific 21103049) supplemented with 1X N2 (Thermo Scientific 17502-048), 0.5X B27 (Thermo Scientific 17504044), 50uM 2-mercaptoethanol, 2mM glutaMAX, 100U/mL penicillin, 100ug/mL streptomycin, 1uM PD032590 (Sigma-Aldrich PZ0162), 3uM CHIR99021 (Sigma-Aldrich SML1046), and 100U/mL LIF (Sigma-Aldrich ESG1106). To introduce plasmid DNA, mESCs were transfected using Lipofectamine 2000 reagent. All transfections were conducted according to the manufacturers protocol with the following parameters per well for a 12-well format: seeding density: 60,000 cells, 0.8ug plasmid DNA, 2.5uL Lipofectamine reagent. For other plate formats, parameters were scaled according to surface area.

### Negative stain electron microscopy

Placentas were harvested and immediately placed in PBS on ice. ~1mm^3^ cubes were cut surrounding the umbilical cord and fixed in Karnovsky’s fixative (2% glutaraldehyde (Electron Microscopy Sciences 16000), 4% PFA (Electron Microscopy Sciences 15700), in 0.1M sodium cacodylate pH7.4 (Electron Microscopy Sciences 12300) for 1h at room temperature. Samples were further trimmed, washed 2X in 0.1M sodium cacodylate, then post-fixed in 2% osmium tetroxide (Electron Microscopy Sciences 19100) for 2h at room temperature. Samples were then incubated in 2.5% v/v potassium ferricyanide (Polysciences 03049) in 0.1M sodium cacodylate for 1.5h, washed 2 x 30m in H_2_O, stained in 1% thiocarbohydrazide (Electron Microscopy Sciences 21900) for 20m, washed 2 x 30m in H_2_O, and *en bloc* stained in 1% uranyl acetate (Electron Microscopy 22400) at 4C overnight. The following morning, samples in uranyl acetate were moved to 50C for 2h, washed 2 x 30min in H_2_O, and *en bloc* stained in Walton’s lead aspartate pH5.1 with potassium hydroxide (Frontier Science 135048) for 2h at 50C. Samples were transferred to lead nitrate (Electron Microscopy Services 1790025), then washed 2 x 30min in H_2_O. Samples were dehydrated via serial ethanol washes (30%, 50%, 70%, 95%, 100%, 100%) for 30min each at room temperature, then submerged in acetonitrile for 30m. Samples were infiltrated with 1:3 Embed-812 resin (Electron Microscopy Sciences 14120) mixed with acetonitrile for 2h, 1:1 Embed-812:acetonitrile for 2 hours, and 3:1 Embed-812:acetonitrile overnight at room temperature. The following morning, samples were placed in Embed-812 for 2 hours, then placed in capsules filled with fresh resin before being incubated at 65C overnight. 75-90nm sections were cut on a Leica UC7 (Leica, Wetzlar, Germany) and mounted onto formvar/carbon coated slot grids (Electron Microscopy Sciences FCF2010-Cu) or 100 mesh copper grids (Electron Microscopy Sciences FCF100-Cu). Grids were imaged on a JEOL JEM-1400 TEM at 120kV and images were acquired with a Gatan Orius digital camera and saved as .dm3.

### Hybridization chain reaction (HCR v3.0)

All HCR probe sets were procured from Molecular Instruments. Target sequences for the probe sets are as follows: IAPEz-int probe set was designed to consensus regions of published IAP sequences from the following publications: ^23,39^. *Iap(fh)* probe set was designed against the IAPE-D1 sequence published and characterized in Ribet, et al. (2008)^24^

Tissues on slides were removed from −80C storage and dried at 37C for 10 minutes. Slides were dehydrated for 5 minutes each through 50% EtOH, 70% EtOH, 100% EtOH (2X). Slides were rinsed in PBS and treated with 10ug/mL proteinase K (Sigma-Aldrich 3115879001) for 5 minutes at room temperature. Slides were washed in PBS again, then prehybridized in prewarmed hybridization buffer for 10 minutes at 37C. Probe sets were diluted to 0.4pmol in hybridization buffer and added to replace prehybridization buffer. Tissues were incubated overnight with probe mixes in a humidified chamber at 37C. The following day, slides were washed for 15 minutes each through graded wash buffer/5x SSCT solutions as follows: 75% wash buffer, 50% wash buffer, 25% wash buffer, 0% wash buffer. Slides were washed with 5 x SSCT 2 x 5 minutes additionally. Amplification buffer was added to slides and incubated for 30 minutes at room temperature. During this time, hairpins were separately snap cooled by heating to 95C for 90 seconds and allowing to cool in the dark for at least 30 minutes. Hairpins were then diluted to 6pmol each in amplification buffer and added to slides. Slides were incubated overnight in a humidified chamber at room temperature protected from light. The following day, slides were washed briefly in 5x SSCT and counterstained with 250ug/mL DAPI (Fisher Scientific EN62248) in 5x SSCT. Slides were further washed 2 x 30 minutes in 5x SSCT before mounting with ProLong Gold (Thermo Scientific P36930). Coverslips were added and slides were allowed to cure for 24 hours before imaging.

For wholemount *in situ*, tissues were removed from cold storage and prehybridized for 5 minutes in prewarmed hybridization buffer. Probe sets were diluted to 2pmol in hybridization and tissues were incubated with probes overnight at 37C. The following day, probes were removed and tissues were washed 4 x 15 minutes in prewarmed wash buffer. During washes, hairpins were snap cooled as above. Embryos were further washed 2 x 5 minutes in 5x SSCT and pre-amplified in amplification buffer for 5 minutes at room temperature. Hairpins were diluted to 30pmol in amplification buffer and embryos were incubated in hairpin mix overnight at room temperature, protected from light. The following day, embryos were briefly washed in 5x SSCT and counterstained with 500ug/mL DAPI in 5x SSCT. Embryos were then washed 2 x 30 minutes in 5x SSCT and stored in 5x SSCT before imaging.

### RNA sequencing

#### Library preparation

Strand specific polyA+ mRNA libraries were generated from 100ng total RNA extract using the NEBNext Ultra Directional RNA Library Prep Kit for Illumina (New England Biolabs E7760), in combination with NEBNext poly(A) mRNA Magnetic Isolation Module (NEB) according to the manufacturers protocol. Library quality and quantity were assessed via Qubit (Invitrogen) and Bioanalyzer (Agilent) before sequencing (2 x 100bp) on an Illumina NextSeq 500.

#### Sample sizes

For E6.5 samples, 10 litters (5 each replicate, corresponding to 27 and 35 conceptuses respectively) were pooled to generate the two replicates. For E9.5 and E15.5 samples, all replicates pooled three independent embryos from three independent dames.

#### Mapping and bioinformatics

Sequences were processed using trimgalore (https://github.com/FelixKrueger/TrimGalore) (adapter trimming, Phred score > 30) and quality controlled using FastQC^64^. Reads were then aligned to the mouse genome (mm39, UCSC) using STAR^33^ (v2.5.3a) with the following parameters:

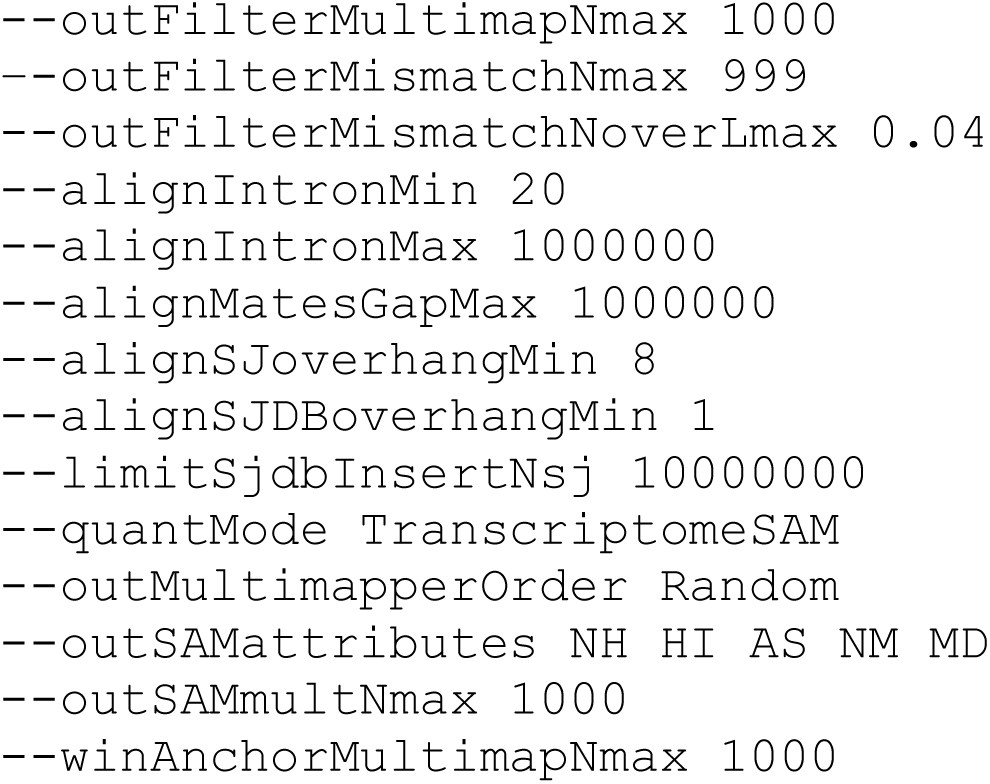

Genes and transposons were downloaded from UCSC tracks (RefSeq and RepeatMasker-viz, respectively). Briefly, for multimapping reads, the primary location was assigned to the best scoring alignment. For cases of equal best scoring alignments, reads were randomly assigned to one of the loci. Transcript expression was quantified with the TEcount module of TEtranscripts^34^ (–mode multi) and normalized using DESeq2^35^.

For all other analyzed species, publicly available RNA-sequencing data sets were utilized (see **Supplemental File 1**). Mapping and bioinformatics were performed as above with the following changes: Phred score > 20, all splice aware mapping parameters were omitted, only repeat elements were counted.

### Statistics and reproducibility

#### Expression outliers

Linear regression was performed for the log_2_(size of family) versus log_2_(normalized counts), where size of family is defined in base pairs as the total length of all elements in the family. Studentized residuals were then calculated for each element and all elements +/− 3 studentized residuals were considered “expressed” or “repressed” respectively. Regression and residual calculations were done in R (v4.3.3).

All other statistical analyses were conducted in GraphPad Prism software (version 10).

### Total protein harvest and extraction

For cell culture supernatants, culture media was removed and immediately placed on ice. Proteins were then concentrated down to ~200uL with an Amicon device with a 10 MWCO cutoff (Millipore Sigma UFC901008). For cell culture lysates, media was removed and cells were rinsed with ice cold PBS. PBS was aspirated and replaced with 1mL of RIPA buffer (150mM NaCl, 50mM Tris-HCl, 1% NP-40, 0.5% sodium deoxycholate, 0.1% SDS, 1 tablet cOmplete protease inhibitor per 50mL). Cells were scraped to disrupt adhesion to the culture plastic. Cells were transferred to an Eppendorf tube and agitated for 2 hours at 4C. Debris was pelleted via centrifugation at 12 000 rpm for 20 minutes at 4C. The resulting supernatant was then transferred to a fresh tube. For whole tissue extraction, tissues were dissected as described above and immediately snap frozen. For protein extraction, tissues were thawed in cold RIPA buffer on ice, then homogenized in 10 second bursts using a rotor/stator. Homogenates were then transferred to Eppendorf tubes and processed as above for cell culture lysates. All total protein samples were stored at −20C prior to use.

### Antibodies

The following primary antibodies were used for western blot: HA (mouse, 1:1000, Abcam ab 9100), GFP (rabbit, 1:2000, Abcam ab13970), CANX (rabbit, 1:1000, Abcam ab213243), CD81 (mouse, 1:500, Abcam ab205606). The following secondary antibodies were used for western blot: anti-mouse HRP (goat, 1:5000, Jackson ImmunoResearch 115-035-003), anti-rabbit HRP (goat, 1:5000, Jackson ImmunoResearch 111-035-003).

### Identification of intact gene domains

To identify putative VLP-competent ERVs, FASTA sequence was extracted for all annotated LTR loci. All ORFs contained within these sequences were extracted, requiring a minimum length of 300bp (100aa). BLAST^37^ and HMMER^38^ were then used to compare ORFs to annotated viral gag domains. Hits passing the following thresholds for both BLAST and HMMER were considered genuine gag ORFs:

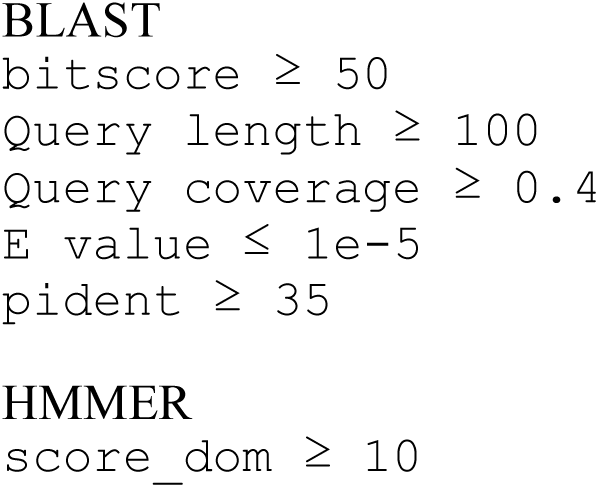

### Copy number estimation

DNA was extracted via chloroform extraction from cell pellets. Sso Universal SYBR mastermix (Bio-Rad 1725272) was used with 150ng input gDNA. Primer sets were constrained to having at least one primer span an intron-exon junction where possible. Primer sequences are as follows (target_species): Gapdh_Hs-F: TCGGAGTCAACGGATTTGGT, Gapdh_Hs-R: TTCCCGTTCTCAGCCTTGAC; Gapdh_Mm-F: GGCAAATTCAACGGCACAGT, Gapdh_Mm-R: CTTTCCGGCCACTTACCCC; EGFP-F: GTTCTGCTGGTAGTGGTCGG, EGFP-R: AGAACGGCATCAAGGTGAACT. Cycling was performed according to manufacturer’s protocol (gDNA input) on a Bio-Rad CFX96 thermocycler. Relative copy number was performed using the ddCt method.

### qPCR

Cell pellets or tissues were harvested and resuspended in Trizol. Total RNA was extracted with the PureLink RNA mini kit (Life Technologies 12185010) according to manufacturers protocol for samples in trizol and treated with DNAse on column to remove contaminating gDNA. First strand cDNA was synthesized with SuperScript III first strand synthesis kit (ThermoFisher Scientific 18080051) with random hexamer primers. 1uL first strand synthesis was mixed with Sso Universal SYBR mastermix. Primer sets were constrained to span an intron to discriminate between any remaining gDNA contamination. Primer sequences are as follows (target_species): Gapdh_Mm-F: ACCCTTAAGAGGGATGCTGC, Gapdh_Mm-R: CCCAATACGGCCAAATCCGT; *Iap(fh)_*Mm-F*: TGCTGGATGTCACAATGCTG, Iap(fh)*_Mm-R: ACCAAAGTCTCGCTTTTGCC; IAPEz-int_Mm-F: AAGGCCAGTTTGCTGATTGG, IAPEz-int_Mm-R: ATGAATGAGTCTGCGCACTG. Cycling was performed according to manufacturer’s protocol (cDNA template) on a BioRad CFX96 thermocycler. Relative expression was calculated using the ddCt method.

### Fluorescence Microscopy and image analysis

Fluorescence images were obtained on a Nikon Ti Eclipse inverted microscope equipped with an ASI MS-2000 motorized linear XY stage, Yokogawa CSU-W1 single disk (50 μM pinhole) spinning disk unit, Andor iXon DU-897 (16 μM pixel size) EMCCD camera, and 10×/0.45 NA Nikon PlanApo Lambda air or 60×/1.4 NA Nikon PlanApo Lambda oil objectives. The final digital resolution of the images was 1.6 μM/pixel for and 0.27 μM/pixel for 10X and 60X images respectively. Blue, red, and far-red fluorescence was collected by illuminating the sample with a 405, 561, or 640nm laser, respectively, in a SPECTRA laser launch, then acquired sequentially through an ET450/40 M (blue), ET595/50 M (red), or ET690/50 M (far red) emission filter. Nikon Elements v4.30.02 was used to acquire the data. Images of entire placental and uterine tissues were captured by tile-scanning the tissue with 15% overlap for stitching after manually identifying XY tissue boundaries. Tiles were manually focused in Z across the scan area to generate a focus surface that determined precise z-position during acquisition. Data were saved in .nd2 format and manually processed in ImageJ^65^. For analysis of HCR images in fetally-restricted *Iap(fh)*, all images were captured with the same exposure settings and intensities were thresholded based on appropriate no amplifier controls.

### Flow cytometry and calculation of transposition frequency

To prepare cell suspensions for flow cytometry, culture media was aspirated and cells were washed with PBS. Cells were then trypsinized for 5 minutes and pelleted after quenching with an equal volume FBS-containing medium. The supernatant was aspirated and cell pellets were resuspended in fresh culture media. Flow cytometry was performed on an Attune Acoustic Focusing Flow Cytometer running v6.2.2 software. Voltages were held constant across experiments and all conditions were run in triplicate. Data were saved in .fcs format and analyzed with FlowJo v10.10.0. Gates were set as follows: FSC-A/SSC-A to determine live/dead cells, SSC-H/SSC-A to isolate singlets, and BL1 to determine GFP positivity. Gates were set on untransfected controls cultured and harvested alongside experimental conditions and applied to all samples per experiment. Transposition frequency was defined as the percentage of GFP expressing cells observed in the (tr)IAP^EGFP^ transfected samples normalized by the transfection efficiency of the experiment (the highest average percentage GFP+ in pEGFP control transfected cells).

## Acknowledgments

The authors thank John Perrino and the TEM team of the Cell Sciences Imaging Facility at Stanford University for their expertise, as well as Hong Zeng and the Tumor, Knockout and Tumor Model Center for their guidance on the generation of the R26-IAP line. The authors would also like to thank Kelly McGowan and Chris Kaelin for reagents and expertise. This work was supported by R01HD110888 (JCB, AJB) and T32HG000044 (AJB). GC was supported by a Walter and Idun Berry Postdoctoral fellowship.

This project was supported in part by NIH S10 Award 1S1OD028536-01 from the Office of Research Infrastructure Programs (ORIP). The contents are solely the responsibility of the authors and do not necessarily represent the official views of the NCRR or the NIH.

## Author Contributions

AJB designed and carried out experiments, data analysis, and visualization. GC designed, performed, and analyzed the RNAseq experiment. JCB supervised all aspects of the project, including design and interpretation. AJB and JCB wrote the manuscript. AJB, GC, and JCB reviewed and edited the manuscript.

