## Supplemental Materials for "Endogenous retrovirus IAP forms virus-like particles and traffics across the maternal-fetal barrier"

**This file includes**
Supplemental Figures 1-2
Supplemental Tables 1-3
Caption for Supplemental File 1

**Other supplementary materials for this manuscript include**
Supplemental File 1 (separate file)

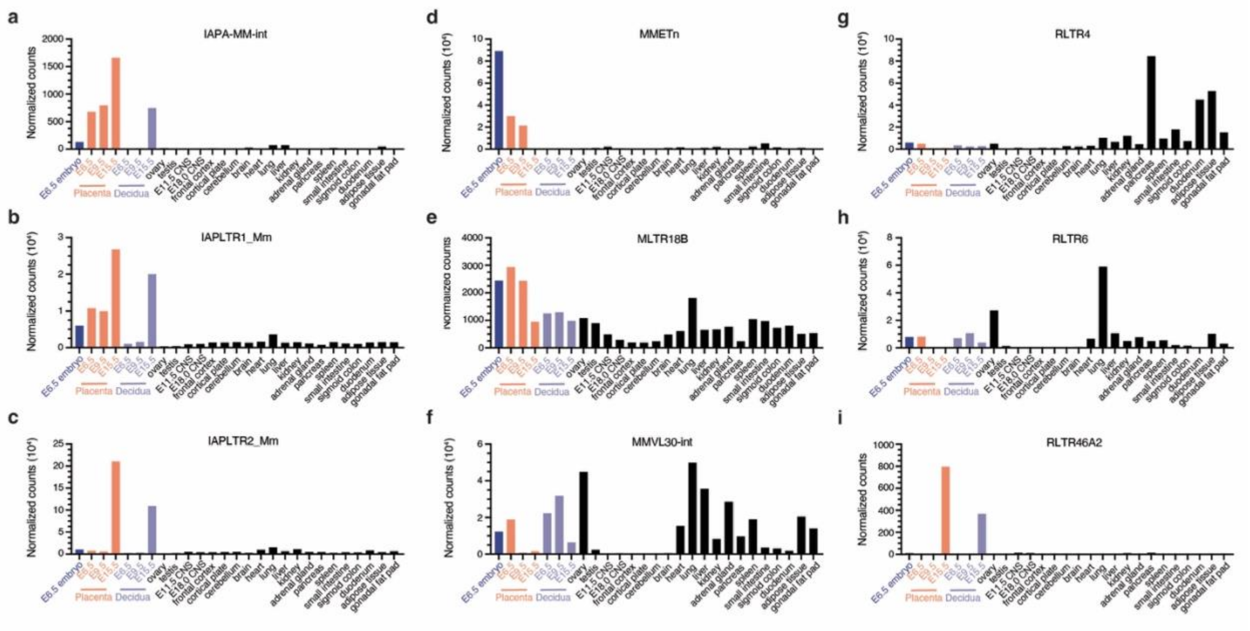


**Supplemental Figure 1 – *Expressed IAP families have unique tissue specificity across gestation***

Expression of identified outlier ERVs in E6.5 embryo (blue), E6.5, E9.5, E15.5 placenta (orange) and decidua (purple), and somatic tissues of the adult and embryo (black). **a)** IAPA-MM-int **b)** IAPLTR1_Mm **c)** IAPLTR2_Mm **d)** MMETn **e)** MLTR18B **f)** MMVL30-int **g)** RLTR4 **h)** RLTR6 **i)** RLTR46A2.

**a**

| Genotype | Observed | Expected |
|---|---|---|
| *Iap(fh) / Iap(fh)* | 17 | 22.5 |
| *Iap(fh) / wt* | 44 | 45 |
| wt / wt | 29 | 22.5 |

p = 0.197, ns

**b**

| Sex | Observed | Expected |
|---|---|---|
| Females | 48 | 45 |
| Males | 42 | 45 |

p = 0.527, ns

**c**

| Dame genotype | Sire genotype | Litter size at birth |
|---|---|---|
| wt / wt | wt / wt | 6.2 +/- 0.2 |
| *Iap(fh) / Iap(fh)* | *Iap(fh) / Iap(fh)* | 6.2 +/- 2.490 |
| *Iap(fh) / wt* | *Iap(fh) / wt* | 6.9 +/- 1.595 |
| wt / wt | *Iap(fh) / wt* | 6.571 +/- 1.618 |

all p > 0.4, ns

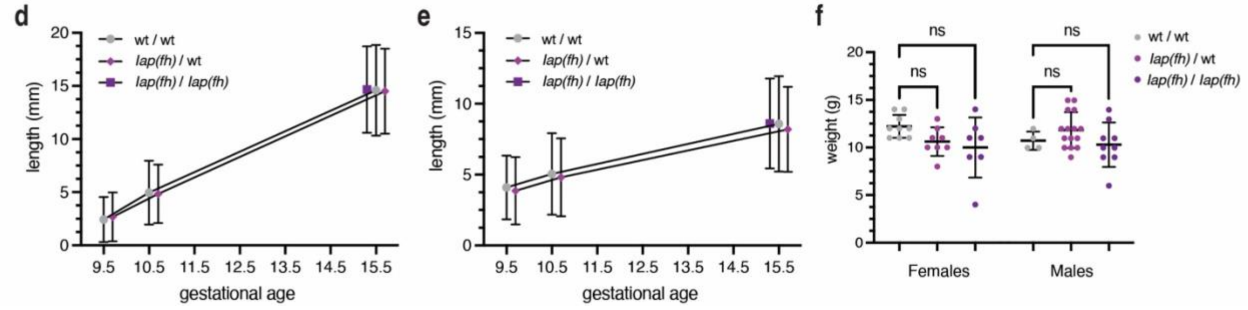


**Supplemental Figure 2 –** ***Iap(fh) does not affect survivorship, fecundity, or growth***

**a)** Chi-square table of genotypes of live offspring from hybrid crosses of R26-IAP parents. Statistics calculated by $\chi$-square test. **b)** Chi-square table of sex of live offspring from hybrid crosses of R26-IAP parents. Statistics calculated by $\chi$-square test. *n=90 animals from 6 crosses.* **c)** Litter size of R26-IAP crosses compared to wildtype (mean, standard deviation). Statistics calculated by Welch's ANOVA with Dunnet's multiple comparisons correction. *n=5-10 litters per class*. **d)** Mean crown-rump length of wildtype (gray), *Iap(fh)* heterozygotes (light purple), and *Iap(fh)* homozygotes (dark purple) embryos across gestation. Error bars represent 95% confidence interval. *n=5-23 embryos from 1-3 litters.* **e)** Mean placental diameter of wildtype (gray), *Iap(fh)* heterozygotes (light purple), and *Iap(fh)* homozygotes (dark purple) embryos across gestation. Bars represent 95% confidence interval. *n=5-23 embryos from 1-3 litters*. **f)** Mean animal weight at weaning (21-25 days post birth) of wildtype (gray), *Iap(fh)* heterozygotes (light purple), and *Iap(fh)* homozygotes (dark purple) animals. Females are plotted in the left panel *(mean weight, by genotype left to right = 12.22, 10.63, 10g)*; males are plotted in the right panel *(mean weight, by genotype left to right = 10.75, 11.87, 10.33g)*. Error bars represent mean and standard deviation. Statistics calculated by one way ANOVA with Dunnet's correction for multiple comparisons. *n=4-9 embryos from 2-5 litters. ns = not significant.*

| E6.5 | E9.5 | E15.5 |
|---|---|---|
| IAPA_MM-int | IAPA_MM-int | IAPA_MM-int |
| IAPLTR1_Mm | IAPEz-int | IAPEz-int |
| MMETn-int | IAPLTR1_Mm | IAPLTR1_Mm |
| MMVL30-int | MLTR18B_MM | IAPLTR2_Mm |
| RLTR4_Mm | MMETn-int | RLTR4_Mm |
| RLTR6_Mm | | RLTR46A2 |

**Supplemental Table 1 –** ***The developing placenta expresses limited ERV families***
Complete list of expressed ERV families in the developing placenta at E6.5, E9.5, and E15.5.

| Family | *gag* | *rt* | *env* | *gag* + *rt* | *gag* + *rt* + *env* |
|---|---|---|---|---|---|
| IAPA_MM-int | 2 | 0 | 0 | | |
| IAPEz-int | 2508 | 1279 | 544 | 1057 | 160 |
| IAPLTR1_Mm | 0 | 0 | 0 | | |
| IAPLTR2_Mm | 0 | 0 | 0 | | |
| MLTR18B_MM | 0 | 0 | 0 | | |
| MMETn-int | 0 | 0 | 0 | | |
| MMVL30-int | 1 | 0 | 0 | | |
| RLTR4_Mm | 0 | 0 | 0 | | |
| RLTR46A2 | 0 | 0 | 0 | | |
| RLTR6_Mm | 0 | 0 | 0 | | |

**Supplemental Table 2 –** ***IAPEz-int loci are uniquely capable of forming VLPs***
Copies containing intact domains with predicted homology among placental-expressed families.

| *Species* (common name) | Expressed families | VLP candidate families |
|---|---|---|
| *Bos taurus* (cow) | LTR6_BT | None |
| *Canis familiaris* (dog) | CarERV3-int<br>CarLTR2<br>MLT1L | None |
| *Dasypus novemcinctus*<br>(9 banded armadillo) | LTR16E1 | None |
| *Equus caballus* (horse) | ERV-I_EC<br>LTR2_EC<br>LTR20_EC | None |
| *Homo sapiens* (human) | ERV3-16A3_I-int<br>ERVL-E-int<br>HERV9NC-int | ERVL-E-int<br>HERV9NC-int<br>HERV9N-int |

| | | |
|---|---|---|
| | HERV9N-int<br>HERVE-int<br>HERVH-int<br>HUERS-P1-int<br>LTR13<br>LTR16A1<br>LTR16B1<br>LTR16<br>LTR24B<br>LTR26E<br>LTR37A<br>LTR37B<br>LTR43<br>LTR43-int<br>LTR78<br>LTR7B<br>LTR8<br>MamGyp-int<br>MER41C<br>MER4A1<br>MER4-int<br>MER52A<br>MER52C<br>MER52D<br>MER57A-int<br>MER68<br>MLT1A0<br>MLT1B<br>MLT1C<br>MLT1C-int<br>MLT1D<br>MLT1E3-int<br>MLT1E<br>MLT1F<br>MLT1G1<br>MLT1G3<br>MLT1H<br>MLT1J2<br>MLT1J<br>MLT1K<br>MLT1L<br>MLT1M<br>MSTA<br>MSTB1<br>MSTB1-int<br>THE1B<br>THE1C-int<br>THE1D | HERVE-int<br>HERVH-int<br>LTR8<br>MER41C<br>MER52A<br>MER57A-int<br>MLT1A0<br>MLT1B<br>MLT1C<br>MLT1D<br>MLT1F<br>MLT1J<br>MSTA<br>MSTB1<br>THE1B<br>THE1D |
| *Loxodonta africana* (elephant) | None | None |
| *Monodelphis domestica* (opossum) | ERV4_MD_I-int | ERV4_MD_I-int |

| | | |
|---|---|---|
| *Mus musculus*<br>(house mouse) | See **Supplemental Table 1** | See **Supplemental Table 2** |
| *Ovis aries* (sheep) | LTR16D1<br>MER21C<br>MLT1B<br>MLT1C2<br>MLT1D<br>MLT1E1<br>MLT1H<br>MLT1K<br>MLT1L | None |
| *Pan paniscus (bonobo)* | MLT1C2 | None |
| *Rattus norvegicus (rat)* | ERVB1_2-LTR_RN<br>MamGyp-int<br>MER34B-int<br>MER92B<br>MLT2F<br>MLTR14<br>MT2B<br>NICER19A-int<br>NICER19B-int<br>NICER2_Rn<br>NICER-int<br>NICER_Rn<br>RAL_Rn<br>RAL_Rn-int<br>RLTR01_Rn<br>RLTR19-int<br>RLTR31_Rn2<br>RMER15<br>RMER2<br>RMER6C<br>RMER6D<br>RNLTR12<br>RNLTR12-int<br>RNLTR1a-int<br>RNLTR3a<br>RNLTR3a-int<br>RNLTR5B<br>RNLTRE<br>RNNICER2-int | NICER19A-int<br>NICER-int<br>RAL_Rn-int<br>RNLTR12-int<br>RNLTR3a-int<br>RNNICER2-int |
| *Macaca mulatta*<br>(rhesus macaque) | HERV4_I-int<br>HERVFH21-int<br>HERVH48-int<br>HERVH-int<br>HERVK-int<br>LTR35<br>MacERV2_int-int<br>MacERV2_LTR2a<br>MacERV3_int-int | HERV4_I-int<br>HERVFH21-int<br>HERVH48-int<br>HERVH-int<br>HERVK-int<br>MacERV2_int-int<br>MacERV3_int-int<br>MacERV5a-int<br>MacERV6-int |

| | | |
|---|---|---|
| | MacERV5a-int<br>MacERV6-int<br>MacNERV5-int<br>MacNERV5_LTR<br>MacNERV6a-int<br>MacNERV6b-int<br>MacNERVK2a-int<br>MER50-int<br>MER65-int<br>MER66B<br>MER66-int<br>MLT-int | MacNERV5-int<br>MacNERV6a-int<br>MacNERV6b-int<br>MacNERVK2a-int<br>MER50-int<br>MER66-int |
| *Sus scrofa* (pig) | ERV1-2B_SSc-LTR<br>ERV1-2_SSc-I<br>ERV1N-1A2_SSc-I<br>MER90a | ERV1-2_SSc-I<br>ERV1N-1A2_SSc-I |

**Supplemental Table 3 –** ***ERVs expression and VLP-competency is pervasive among mammalian species***
Expressed and candidate VLP families identified in all species analyzed.

**Supplemental File 1 (separate)**
Data used for analysis of ERV expression and VLP candidate search in all species. Data sets were pooled as replicates by gestational age (as a percentage of total gestation length, rounded to the nearest 10%). All numbers are reported first by age, then by total summed across all gestational ages. Number of VLP candidate families refers to the number of expressed families containing at least one locus with a VLP candidate locus. Number of VLP candidate loci refers to the number of VLP-competent loci across all expressed ERV families.
